# Tutorial: guidelines for manual cell type annotation of single-cell multi-omics datasets using interactive software

**DOI:** 10.1101/2023.07.11.548639

**Authors:** Yang-Joon Kim, Alexander Tarashansky, Karen Liang, Meg Urisko, Leah Dorman, Michael Borja, Norma Neff, Angela Oliveira Pisco, Alejandro Granados

## Abstract

Assigning cell identity to clusters of single cells is an essential step towards extracting biological insights from many genomics datasets. Although annotation workflows for datasets built with a *single* modality are well established, limitations exist in annotating cell types in datasets with *multiple* modalities due to the need for a framework to exploit them jointly. While, in principle, different modalities could convey complementary information about cell identity, it is unclear to what extent they can be combined to improve the accuracy and resolution of cell type annotations.

Here, we present a conceptual framework to examine and jointly interrogate distinct modalities to identify cell types. We integrated our framework into a series of vignettes, using immune cells as a well-studied example, and demonstrate cell type annotation workflows ranging from using single-cell RNA-seq datasets alone, to using multiple modalities such as single-cell Multiome (RNA and chromatin accessibility), CITE-seq (RNA and surface proteins). In some cases, one or other single modality is superior to the other for identification of specific cell types, in others combining the two modalities improves resolution and the ability to identify finer subpopulations. Finally, we use interactive software from CZ CELLxGENE community tools to visualize and integrate histological and spatial transcriptomic data.

## 1. Introduction

Advancements in single-cell genomics have revolutionized our ability to examine characteristics of heterogeneous cell populations with remarkable precision (Nomura 2021; Gawad, Koh, and Quake 2016; Sandberg 2014). Initially, single-cell RNA-sequencing (scRNA-seq) enabled the investigation of transcriptomic states of individual cells by measuring RNA molecules at the whole transcriptome level (Macosko et al. 2015; Klein et al. 2015). This technology has facilitated the identification, classification, and examination of most cell populations in various organisms, including developmental and disease states, based on their differences in gene expression (Tabula Muris Consortium et al. 2018; Lange et al. 2023; Saunders et al. 2022; Sur et al. 2023; Delorey et al. 2021; Smillie et al. 2019; The COVID Tissue Atlas Consortium et al. 2022).

However, cellular identity is multifaceted and involves multiple regulatory aspects across biological processes, including chromatin accessibility at the genome level, gene expression profile at the transcriptome level, and protein abundance at the proteome level (Figure 1a and b). Measuring RNA levels *in situ* via spatial transcriptomics provides orthogonal and valuable information as cell identity is defined in the context of tissue organization and cell-cell interactions. Access to these multiple layers of molecular states in single cells and tissues should increase the resolution and accuracy of cell type annotations (Figure 1c), and enable the discovery of new cell populations.

**Figure 1.**
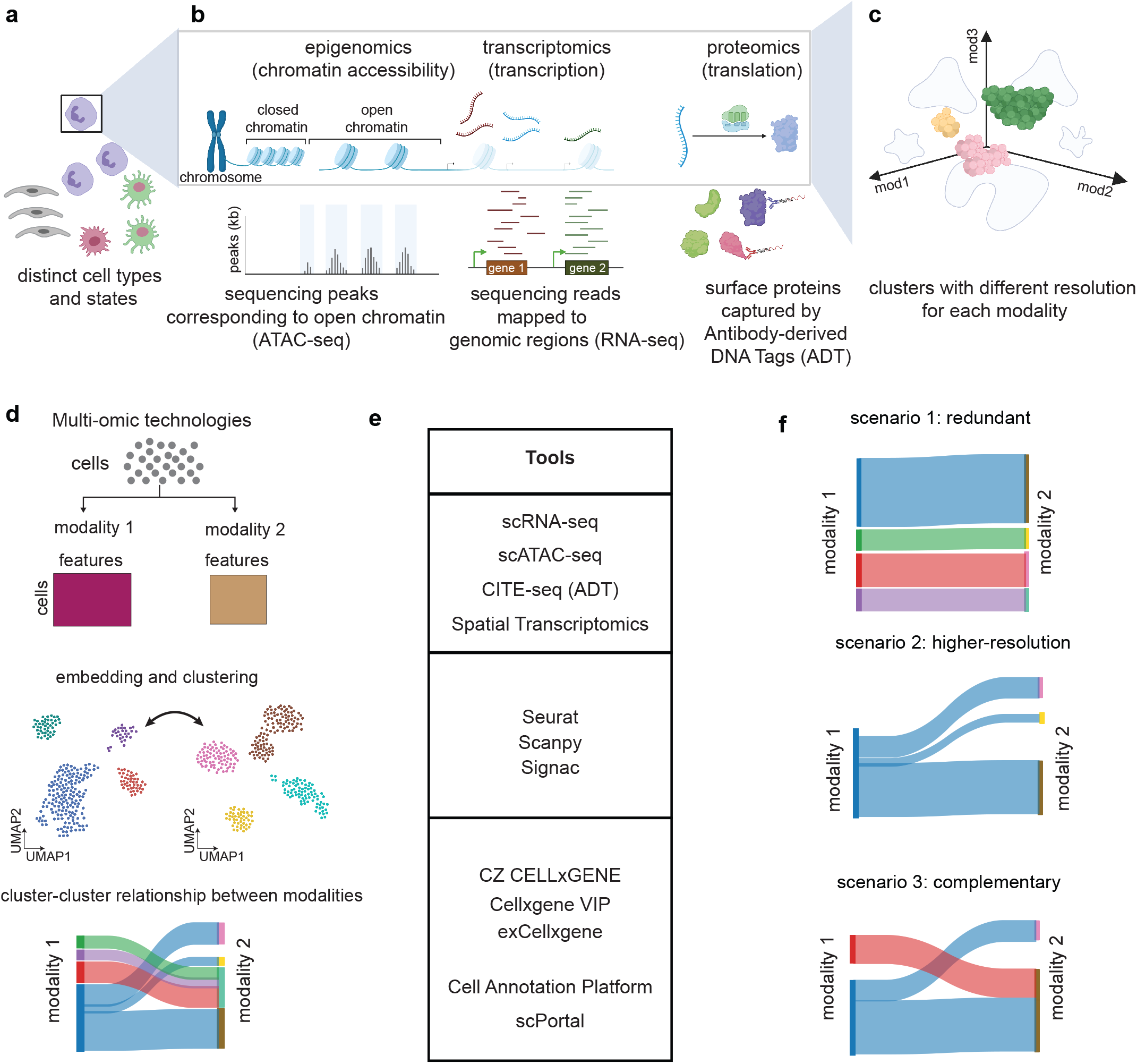
A conceptual framework for cell type annotation in multi-omics datasets. **a)** A schematic showing distinct cell types and states. **b)** A schematic showing different layers of gene expression regulation (modalities) along the Central Dogma (chromatin accessibility, transcription, and translation) and example assays to measure these modalities. **c)** A conceptual schematic showing the resolution of cell types in multi-omics datasets. Each axis represents the aforementioned “modality”, and cell identity can be abstracted as coordinates in 3D modality space. 2D projections represent the clusters of cells resolved by two modalities. **d)** A workflow of multi-omics data exploration: (top) two count matrices generated from multi-omics datasets: cells by features1 (modality1) and cells by features2 (modality2), where the cells are shared between the two modalities. (middle) Each modality is processed independently (Quality Control, dimensionality reduction, clustering, 2D embedding, etc.) giving rise to two 2D embeddings from each modality. (bottom) cluster-cluster relationships between the two modalities examined by interactive software tools, illustrated by a Sankey diagram. **e)** A summary of the tools/packages used for each step in (d). **f)** Different scenarios for the cluster-cluster relationships between the two modalities. (top) *redundant*, the clustering from the two modalities are 1:1 match, giving redundant information about the resolution of cell types. (middle) *higher-resolution*, one modality gives a higher resolution of clusters than the other. (bottom) *complementary*, the resolution of cell types is complementary, meaning that some cell types (clusters) can be revealed by utilizing both modalities.

In recent years, new single-cell multi-omics technologies have emerged that allow for the simultaneous measurement across multiple regulatory regimes including RNA, chromatin accessibility, and protein (Figure 1b) (Ma et al. 2020; W. Xu et al. 2022; Stoeckius et al. 2017; Moses and Pachter 2022b). As a result, accurate annotation of cell populations should encompass all aspects of this multi-dimensional feature space that defines the state of a cell (Figure 1c). While various workflows for cell clustering and annotation for single-cell transcriptomics datasets exist (Luecken and Theis 2019; Clarke et al. 2021), there is a need for standard practices when annotating cell types from multi-omics datasets based on their multi-dimensional molecular states.

Single-cell multi-omics techniques usually measure two different aspects of cell identity simultaneously. Although the specifics of each technology may differ, they essentially provide paired measurements for each cell across two sets of features (Figure 1d - top). Each expression matrix, representing cells-by-features, can be analyzed independently by applying feature extraction, dimensionality reduction, and clustering (Luecken and Theis 2019). This results in two independent two-dimensional (2D) embeddings of the same set of cells (Figure 1d, center). However, the resolution of cellular groupings may differ between modalities since each provides complementary information about the cellular states (Figure 1d, bottom). For example, some cell types can be better resolved in the protein feature space than RNA feature space. In other cases, chromatin accessibility can be used to define fine subtypes of cells whose transcriptional profiles are not yet distinct but whose chromatin accessibilities have been differentiated (Itokawa et al. 2022; Ma et al. 2020). Alternatively, modalities that are highly correlated for specific cell populations may provide redundant information.

The extent to which different modalities provide complementary or redundant information that can be used to annotate cell types is largely unknown (Miao et al. 2021). Although the technologies for generating multimodal datasets are now broadly available, analyzing these datasets often requires proficiency in programming languages such as Python or R (Figure 1e) (Wolf, Angerer, and Theis 2018; Satija et al. 2015; Stuart et al. 2019, 2021). These challenges can be overcome by *interactive* software that democratizes the analysis of multimodal datasets, leveraging the diverse knowledge base of researchers without formal training in programming languages (Figure 1f).

In this tutorial, we present a guideline for exploring and annotating single-cell multi-omics datasets using interactive tools. While there are several interactive tools available for exploring single-cell datasets, most are focused on the RNA modality, and only a few support multimodal datasets. We use a suite of interactive CZ CELLxGENE tools, including exCELLxGENE and Cellxgene VIP, designed for visualizing, exploring, and annotating single-cell transcriptomics datasets (Megill et al. 2021; K. Li et al. 2020). These tools allow users to load their datasets, explore gene expression patterns and annotate cells using a lasso tool (Megill et al. 2021; K. Li et al. 2020).

We provide four vignettes with step-by-step instructions, code, and documentation for annotating cell types. First, we start with a scRNA-seq dataset, then we expand to multi-omics datasets such as CITE seq (RNA + ADT) and single-cell Multiome (RNA + ATAC). We provide examples in which different modalities offer different depths of information regarding cell identities and discuss best practices for identifying novel populations. Finally, we demonstrate how Cellxgene VIP can be used to integrate histologically-stained tissue samples with spatial transcriptomic data obtained using the 10X Visium (RNA + spatial) system.

## Results

### 2. Cell type annotation of a scRNA-seq dataset using interactive features

Automated cell type annotation has become increasingly common in scRNA-seq data analysis workflows due to its fast turn-around time and scalability; however, manual curation by experts is still required (C. Xu et al. 2021; Domínguez Conde et al. 2022; Ji et al. 2023; Pasquini et al. 2021; Clarke et al. 2021). The usual workflow for manual cell type annotation involves unsupervised clustering of dimensionality-reduced data based on transcriptional similarity, followed by tuning clustering resolution and manually checking marker genes. This is, however, often challenging because no single resolution parameter works across all cell types, and cell annotation typically requires collaboration between multiple researchers followed by collation of annotations. Therefore, efficient cell type annotation requires a platform that does not depend on specific programming languages or data formats, while enabling users to perform at least three key tasks: 1) to define clusters based on expression levels of a marker gene or combinations of marker genes, 2) to explore the data at finer resolution by subsetting and reembedding (Figure 2a), and 3) to switch the embeddings between different modalities for a comprehensive analysis of cell identity within each context. These three functionalities are the key features of exCELLxGENE (https://github.com/czbiohub-sf/excellxgene), an interactive tool for single-cell sequencing analysis and visualization developed by the Chan Zuckerberg Biohub - San Francisco that extends the functionality of CZ CELLxGENE Annotate (Box 1) (Megill et al. 2021).

**Figure 2.**
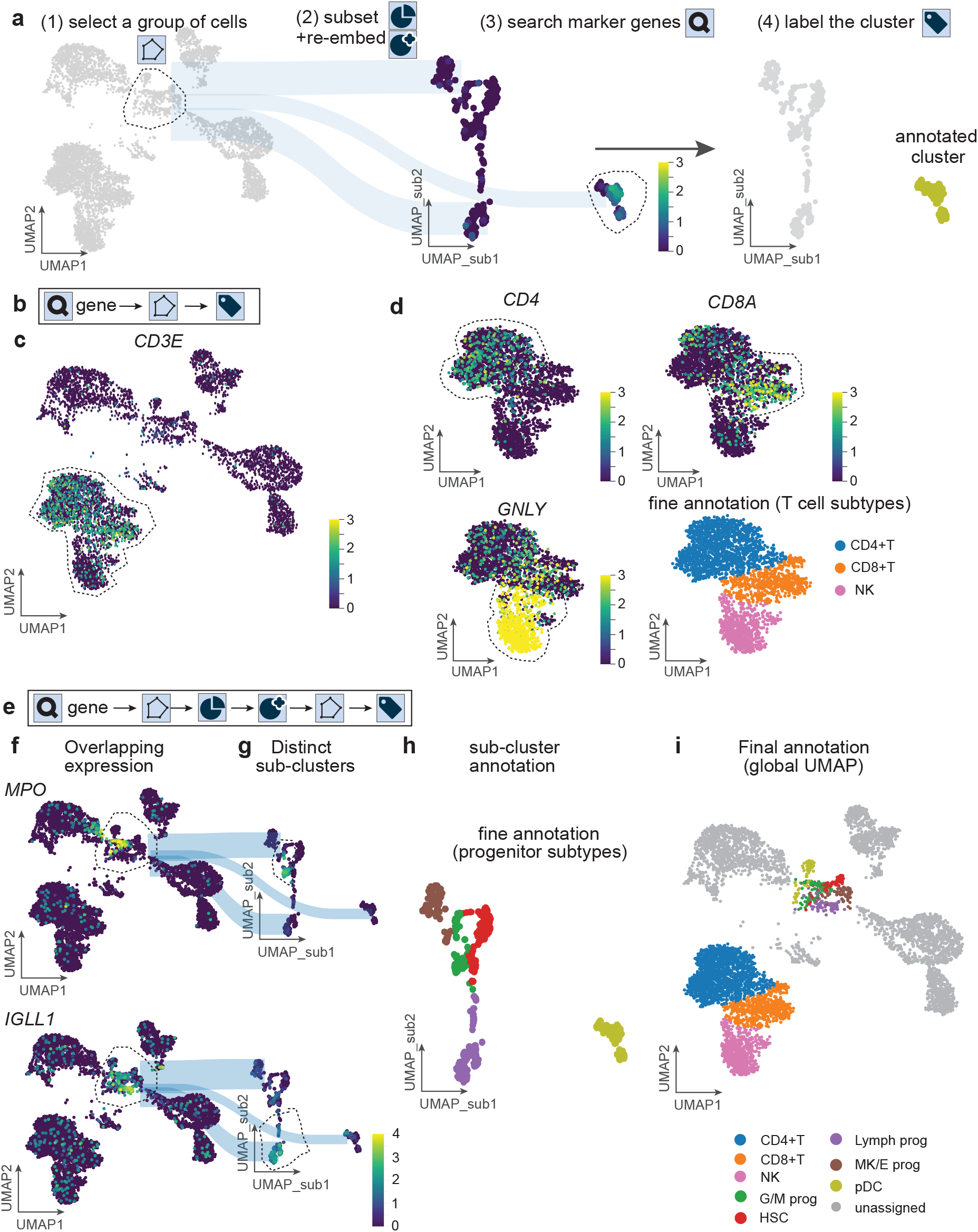
Annotation workflow for single-cell (nucleus) RNA-sequencing dataset with known cell types and marker genes corresponding to each cell type. **a)** Summary of features and corresponding workflow for easier cell type annotation in exCELLxGENE (icons correspond to action buttons on the user interface). **b)** A basic workflow of cell type annotation in exCellxgene using marker genes, as depicted by the action icons shown in (a). **c)** An example of cell type annotation for T cells using *CD3E* as a marker gene following the workflow in (b), using the action icons introduced in (a). **d)** Annotation of T cell subtypes using the subtype marker genes. **e)** An advanced workflow of cell type annotation for populations with overlapping gene expression from multiple subtypes. **f)** A global UMAP showing overlapping marker expression. **g)** subsetted/re-embedded progenitor cells within the lasso in (f). **h)** Fine annotation of progenitor cell types on subsetted/re-embedded UMAP. **i)** Representation of fine-annotated cell types on the global UMAP. G/M prog: Granulocyte/Monocyte progenitors, HSC: Hematopoietic Stem Cells, Lymph prog: Lymphocyte progenitors, MK/E prog: Megakaryocyte/Erythroid progenitors, pDC: plasmacytoid Dendritic Cells.

For this and the following vignettes, we used public datasets from Bone Marrow Mononuclear cells (BMMC). We focus on immune cell subtypes because their markers are well-established and have been used successfully to identify distinct populations in blood and bone marrow. In our annotation workflow, we start with a pre-computed UMAP, and a list of canonical marker genes from the literature (Figure 2a) (Luecken et al. 2022). In principle, a simplified workflow to annotate a cell type within a scRNA-seq dataset would consist of three key steps (Figure 2b, icons correspond to actions in exCELLxGENE interface): 1) Visual inspection of known marker genes of interest in the 2D embedding, 2) Identification of a cluster showing enrichment of the markers, and 3) Labeling of the cluster according to the cell type associated with those markers. For example, a cluster localized at the bottom of the UMAP shows high expression of *CD3E*, a marker gene for T cells (Figure 2c) (DeJarnette et al. 1998). If high-resolution marker genes for subtypes are available, one can identify such populations by looking at the expression of these genes within the cluster in the 2D embedding (Figure 2D). For example, T cell subtype marker genes such as *CD4* (CD4+ T cells), *CD8A* (CD8+ T cells), and *GNLY* (Natural Killer T cells) mark subtypes of T cells based on the surface proteins these genes encode (Figure 2d) (Maecker, McCoy, and Nussenblatt 2012).

Dimensionality reduction algorithms, such as UMAP, face challenges in projecting cell transcriptomes into 2D space while preserving global and local relationships embedded within the underlying data, which can obscure the detection of certain cell types (Becht et al. 2018; Kobak and Berens 2019). This is particularly problematic when annotating small populations with similar transcriptional profiles, such as immune progenitor cells. In these cases, subsetting the data for the specific population and re-computing the embedding for the subset (re-embedding) can help to resolve local structures of neighborhoods for such populations. To address this, the annotation workflow includes additional steps (Figure 2e): 1) Visual inspection of known marker genes of interest in the 2D embedding, 2) Identification of a cluster showing overlapping enrichment of marker genes from several cell types, 3) Subsetting and re-embedding the cluster to reveal local structures, 4) Visual inspection of known marker genes again in the newly generated 2D embedding, and 5) Labeling of the new clusters according to the cell types associated with those markers.

For example, the immune progenitor cells show an overlapping expression of multiple subtype markers, likely because of their potential to differentiate (Figure 2f) (Zhang et al. 2014).

However, their exact identity depends on the expression of specific subtype markers. Granulocyte/Macrophage progenitors (G/M prog) express *MPO*, while Lymphocyte progenitors (Lymph prog) express *IGLL1*, but their expression overlaps in the UMAP (Figure 2f). In this example, subsetting and re-embedding the progenitor populations revealed a more detailed structure in the subset UMAP, where expression of the marker genes further separated when compared to the global UMAP (Figure 2g,h) revealing local structures that would be otherwise obscured by larger global cell differences (Figure 2I).

In summary, we showcased a workflow to annotate cell types using interactive software for a single modality in scRNA-seq datasets. In the following sections, we will explore how to annotate datasets with multiple modalities, namely CITE-seq (RNA+ADT) and single-cell Multiome (RNA+ATAC).

### 3. Annotation of the CITE-seq dataset shows the complementary power of RNA and ADT modalities in resolving finer cell subtypes

While single-cell RNA seq has revolutionized how we understand cell identity, the correlation between cellular RNA levels and their corresponding proteins can vary across cell types, tissues, organisms, and conditions (Reimegård et al. 2021). The CITE-seq assay simultaneously measures gene expression (RNA) and surface protein abundance as targeted by Antibody-derived DNA Tags (ADT) for individual cells (Stoeckius et al. 2017) (Figure 3a). Notably, while the RNA feature space covers the whole transcriptome, the protein feature space, which focuses only on surface-expressed proteins and depends on the size of the ADT panel spanning from tens to hundreds (Nettersheim et al. 2022), provides only a targeted subset of the proteome. After pre-processing, CITE-seq data can be represented as two matrices: a cell-by-gene matrix for RNA and a cell-by-protein matrix for ADT (Figure 3a). For each modality, dimensionality reduction and clustering can be performed separately, resulting in two independent UMAP embeddings (Figure 3a).

**Figure 3.**
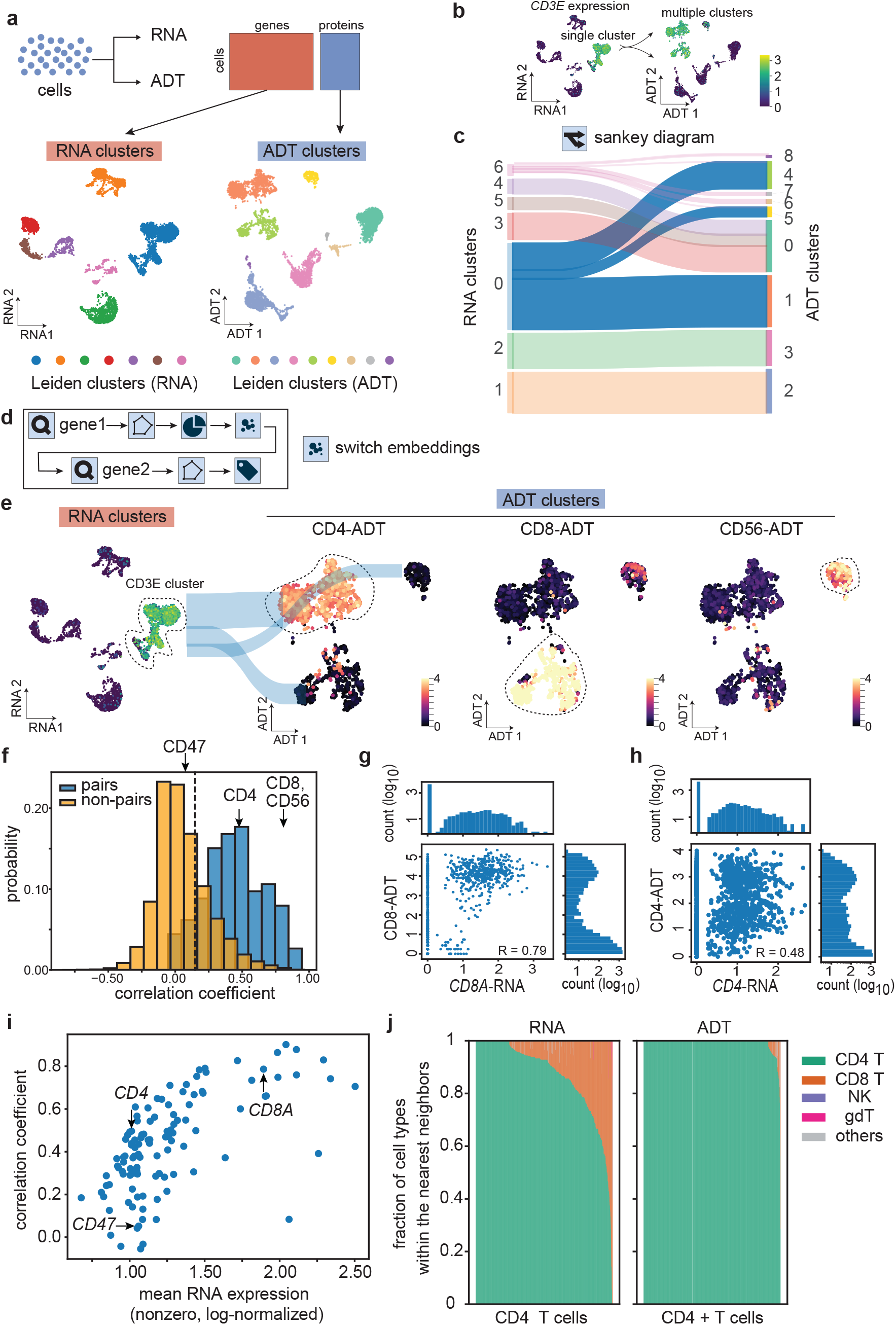
Annotation of a BMMC CITE-seq dataset using both RNA and ADT modalities. **a)** (Top) A schematic for CITE-seq data format. (Bottom) UMAP visualization of each modality with Leiden clustering (resolution = 0.5) **b)** RNA expression for a marker gene for the T cell population, *CD3E*, in RNA and ADT UMAPs. **c)** Sankey diagram between the RNA clusters and ADT clusters shown in (a). Clusters representing T cells are highlighted with the same colors as in (a) **d)** A workflow for cell type annotation from multi-omics datasets using dynamic switching between embeddings from each modality. **e)** An RNA cluster representing the T cell population (left), and zoomed-in ADT UMAP for the T cell population colored with subtype markers. Cells with high expression of each ADT marker are encircled by a black dotted line for annotation and labeling. **f)** A histogram showing the Pearson correlation between RNA and ADT expression at the single-cell level. Histogram consists of 113 pairs of either RNA:ADT pairs (blue) or all non-specific pairs (yellow) to show the background distribution. The dotted line represents the 75 percentile of the distribution from non-specific pairs. **g**,**h)** Scatter plots for RNA (x-axis) and ADT (y-axis) expression for (g) a pair of *CD8A*:CD8, and (h) a pair of *CD4*:CD4, where each dot represents a single cell. Pearson correlation coefficients are shown at the bottom right of the scatter plots. Histograms for each modality (RNA or ADT) are also shown at the top and right of each scatter plot. **i)** A scatter plot for the mean levels of RNA expression across single cells and the Pearson correlation coefficients for each RNA:ADT pair. **j)** Fraction of cell types for each cell (bar) within the nearest neighbors for CD4 T cells for RNA neighborhood (Left) and ADT neighborhood (Right).

In principle, quantifying RNA and protein abundance should increase the information available to classify and label cell populations (Hao et al. 2021). In this section, we used a CITE-seq BMMC dataset that captures immune cell populations, as described in the previous section (Luecken et al., 2022). For example, ADT could provide additional information to split an RNA cluster into three distinct clusters (Figure 3b, *CD3E* gene expression for T cells). To understand whether the ADT panel provides complementary or redundant information relative to RNA about cell identity, we independently analyzed and computed the Leiden clusters for RNA and ADT using the same resolution parameters (Figure 3a, color labels). We then compared the Leiden clusters from the two modalities using the Sankey diagram functionality of exCELLxGENE. A Sankey diagram represents the flow of data from one category to another. In the context of CITE-seq, the Sankey diagram shows the similarities and differences between the two modalities by connecting clusters using flow lines and visualizing the proportion of data points (cells) within each cluster (Figure 3c). In this section, we will focus on RNA cluster 0, which represents the T cell population, that maps to three ADT clusters 1, 4, and 5, to see how we can utilize multiple modalities to define fine subtypes of T cells easily.

Using exCELLxGENE’s switch-embedding feature(Figure 3d) along with the Sankey diagram functionality, we observed that RNA cluster 0 (blue) split into three clusters in the ADT embedding (1,4,5), suggesting that ADT data might achieve a higher resolution in identifying these cell populations (Figure 3c). The top differentially expressed genes were used to identify the RNA cluster 0 as T cells due to the high expression of *CD3E* (Figure 3b). To assess whether the three ADT subclusters represent distinct T cell sub-populations, we analyzed the ADT signal for marker genes associated with T cell subtypes (Figure 2d). Each cluster was enriched for a different marker gene, supporting our observation that ADT offers higher resolution to distinguish T cell subtypes than RNA (Figure 3e, lasso tool was used to highlight the enrichment of each antibody).

While the ADT feature space provided a higher resolution than the RNA data for annotating T cell subtypes, the opposite was true for annotating the erythroblast population. In this case, three RNA clusters (3,4,5) mapped to one ADT cluster (0) (Figure 3c), which we identified as different erythroblast subtypes, proerythroblasts, erythroblasts, and reticulocytes. Therefore, RNA and ADT modalities complement each other in resolving cellular heterogeneities within the immune cell populations.

Differences in RNA and protein expression levels can explain the complementary nature of the RNA and ADT modalities. To assess the contrast between the two modalities, we computed the Pearson correlation between RNA and ADT expression across single cells for each of the 113 RNA:ADT pairs, where each RNA(gene) encodes the corresponding ADT (protein) (Figure 3f, blue). 39 genes showed relatively high correlations (Pearson coefficients > 0.5), including *CD8A*:CD8 (Figure 3g, Pearson coefficient = 0.79); however, the majority of genes showed poor correlations (Pearson coefficient< 0.5), including *CD4*:CD4 (Figure 3h, Pearson coefficient = 0.48). Pairs with high correlation are redundant since we expect them to provide only a marginal amount of extra information about cell identity. In contrast, pairs with low correlation are expected to provide more information than either modality for these genes.

To establish a baseline for noise, we computed the correlation for all non-pairs of RNA and ADT molecules, which corresponds to a null distribution, and compared it with the distribution from the correct RNA:ADT pairs (Figure 3f, yellow). 12/113 pairs showed low correlation coefficients (below the 75th percentile in the null distribution, shown in Figure 3f, dotted line), indicating that RNA and ADT measurements together provide complementary information that can improve resolution (e.g., CD47). Next, we looked for any trend in RNA:ADT pairs’ correlation.

Interestingly, the correlation coefficients tend to increase as the expression levels of RNA increase (Figure 3I). One explanation is that the higher the RNA expression, the easier it is to distinguish the cells expressing the corresponding gene from those showing only noise-level expression. Thus, the signal-to-noise ratio of both RNA and ADT signals could impact the correlation of RNA:ADT expression.

To further explore the accuracy of cell type annotations within a modality, we quantified the fraction of different cell type labels within the nearest neighborhoods of individual cells. For a cell type, if most of the nearest neighbors are from the same cell type, then the cell type is very well separated from the others. As an example, we computed the fraction of cell types within the neighborhoods of CD4 T cells, which showed a better separation in 2D embeddings in ADT than RNA. We used a stacked bar plot to visualize the composition of cell types within the nearest neighbor for each CD4 T cell for RNA and ADT, respectively (Figure 3j). We found that the fraction of homogenous cell type labels (CD4 T cells) is significantly higher in ADT than in RNA. Together, these examples show the power of interactive tools such as exCELLxGENE for accurate annotation of cell types in multi-omic datasets.

## 4. Annotation of Multiome (scRNA+scATAC) dataset shows the complementary power of RNA and ATAC modalities in resolving finer cell subtypes

Transcription factors enable gene activation by binding to regulatory DNA regions and recruiting transcriptional machinery, which in turn initiates transcription (Figure 1a) (He et al. 2013; Nogales, Louder, and He 2017; Levine 2010). ATAC-seq (Assay for Transposase-Accessible Chromatin with sequencing) enables the genome-wide identification of DNA regions accessible to transcription factors (Buenrostro et al. 2013; Yan et al. 2020). Single-cell ATAC-seq

(scATAC-seq) reveals heterogeneity across cell populations based on differential chromatin accessibilities, and it can be used to reconstruct the continuum of cellular differentiation for different cell types during the development of multicellular organisms (Buenrostro et al. 2015; Cusanovich et al. 2015; Cusanovich, Reddington, et al. 2018; Cusanovich, Hill, et al. 2018; Domcke et al. 2020).

In this section, we continued our exercise using the same dataset introduced in Section 2, which encompasses both RNA and ATAC measurements obtained from the identical group of cells (Luecken et al. 2022). To understand the extent to which these different modalities complement each other for identifying cell types, we processed each modality independently and compared the resulting clusters (Figure 4a). In addition to the individual UMAPs, we computed the joint embedding using Seurat v4 (Figure 4b), which applies the weighted nearest-neighbor graph approach using both genes and peaks (Hao et al. 2021). From these three embeddings, we generated a Sankey diagram to compare the Leiden clusters from each modality and the joint embedding (Figure S1). Multiple clusters showed a 1:1 agreement between RNA and ATAC modalities (Figure S1), suggesting that their cell identities can be, in principle, resolved using either modality. However, we noticed cases in which two modalities defined clusters in a complementary fashion. For example, we found a non-1:1 mapping between RNA clusters 5 and 9 and ATAC clusters 7 and 9 (Figure 4c). Although there are two clusters for each modality, three distinct populations emerged when visualizing the flows between RNA and ATAC modalities. We computed the Leiden clusters from the joint embedding, which apparently resolved these three populations. These three Leiden clusters, which can only be identified in the joint embedding (Figure 4c), correspond to the ground-truth annotations for three B cell subtypes (i.e. B1 B cells, Naive CD20+ B cells, and Transitional B cells; Figure SI) with 93.5% match.

**Figure 4.**
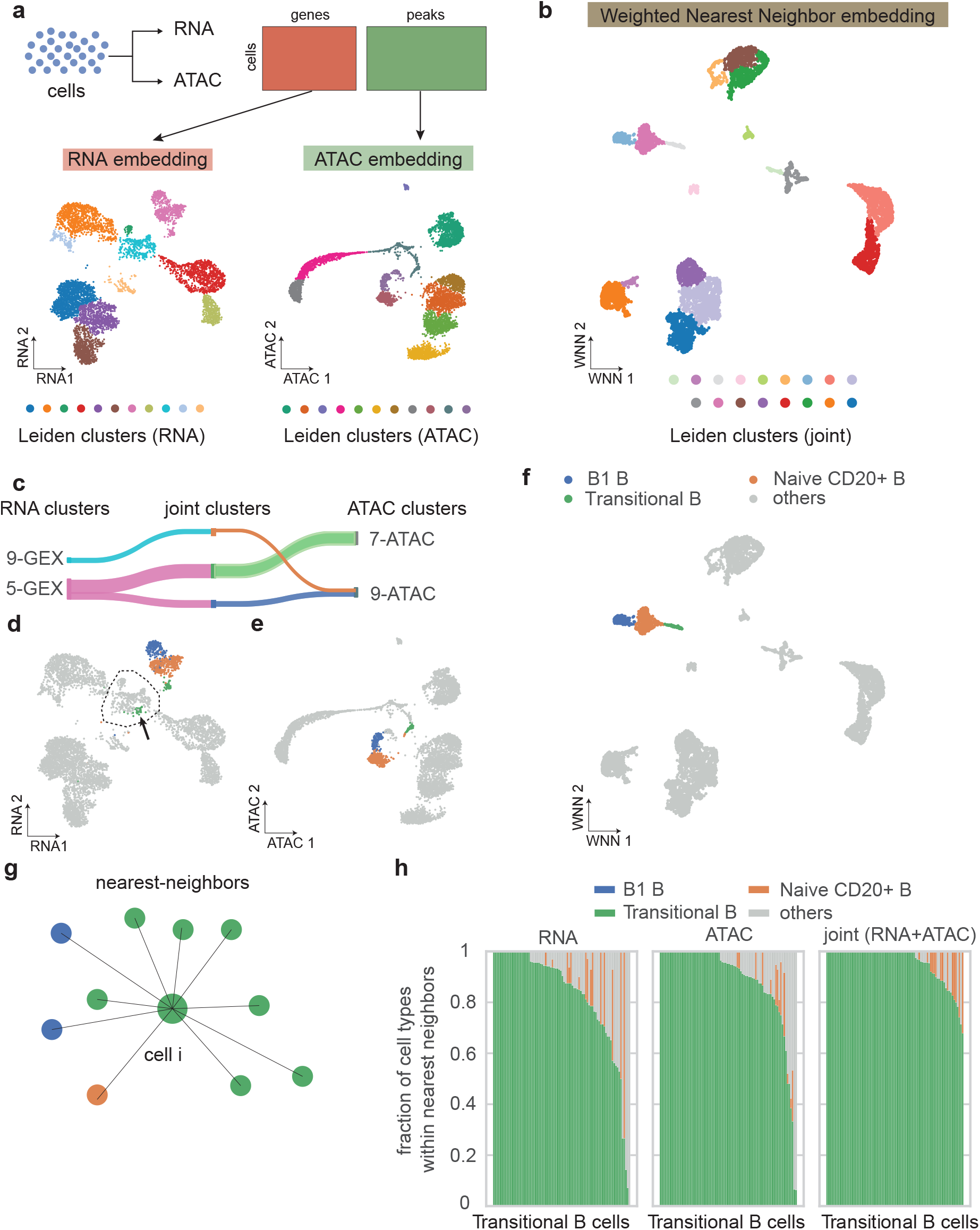
Annotation of a BMMC single-cell multiome dataset (RNA for gene expression and ATAC for chromatin accessibility). **a)** (Top) A schematic for single-cell multiome (scRNA-seq and scATAC-seq) data format. (Bottom) UMAP visualization of each modality with Leiden clustering. **b)** joint embedding (UMAP) from Seurat v4, which applies the weighted nearest neighbor graph approach using both genes and peaks. **c)** Sankey diagram between the RNA clusters and ATAC clusters shown in (a). Only clusters representing B cell subtypes are shown here. **d)** RNA UMAP colored by three subtypes of B cells. The center region outlined by a dotted line represents the immune progenitor cells introduced in Section 2. **e)** ATAC UMAP colored by three subtypes of B cells. **f)** joint UMAP colored by three subtypes of B cells. **g)** A schematic showing the cell type composition in the nearest neighbors of cell i. **h)** Fraction of cell types for each cell (bar) within the nearest neighbors for Transitional B cells in RNA (left), ATAC (center), and joint (right) embeddings.

We projected these three subtypes back into the individual embeddings (RNA or ATAC, Figures 4d and e, respectively) and found that the RNA embedding failed to split Transitional B cells from the immune progenitor cells (highlighted by a dotted line in Figure 4d, annotated in Section 2). Conversely, Naive CD20+ B cells appeared scattered in the ATAC UMAP. However, in the joint embedding, all the B cell subtypes clustered into distinct regions of the UMAP (Figure 4f). To quantify the accuracy in cell annotations in Transitional B cells in each modality, we computed the fraction of cells in single-cell neighborhoods corresponding to each cell type annotation (Figure 4g). We found that Transitional B cells show the highest homogeneity in the nearest neighbors in the joint embedding compared to individual modalities (Figure 4h). In contrast, the RNA embedding showed the highest heterogeneity in the neighborhoods, mostly due to confusion with progenitor cells, suggesting that the Transitional B cells’ chromatin accessibility could be already differentiated from the progenitor cells, but their gene expression profiles have yet to be differentiated. Therefore, by combining ATAC with RNA modalities, we were able to resolve the subtype populations that would have been obscured in the individual modalities.

## 5. Annotation of Spatial Transcriptomics dataset guided by the tissue histology image reveals region-specific pathways

Recently, several spatial transcriptomics technologies have emerged, enabling researchers to investigate cell-cell communication, ligand-receptor interactions, and other questions that require the spatial information of cells within the tissue context (Lein, Borm, and Linnarsson 2017; Longo et al. 2021; Burgess 2019; Rao et al. 2021; Moses and Pachter 2022c). Most spatial transcriptomics technologies accompany imaging data from the same tissue, such as H&E images or immunofluorescence images for nuclei and cell membranes. For tissues where spatial regions are visually identifiable based on cellular morphology, biologists can start cell/tissue annotation directly from the images, without clustering based on transcriptomes (Figure 5a). This advantage offers an orthogonal way of annotating the cells, which can then be verified by an exploration of the gene expression profiles.

**Figure 5.**
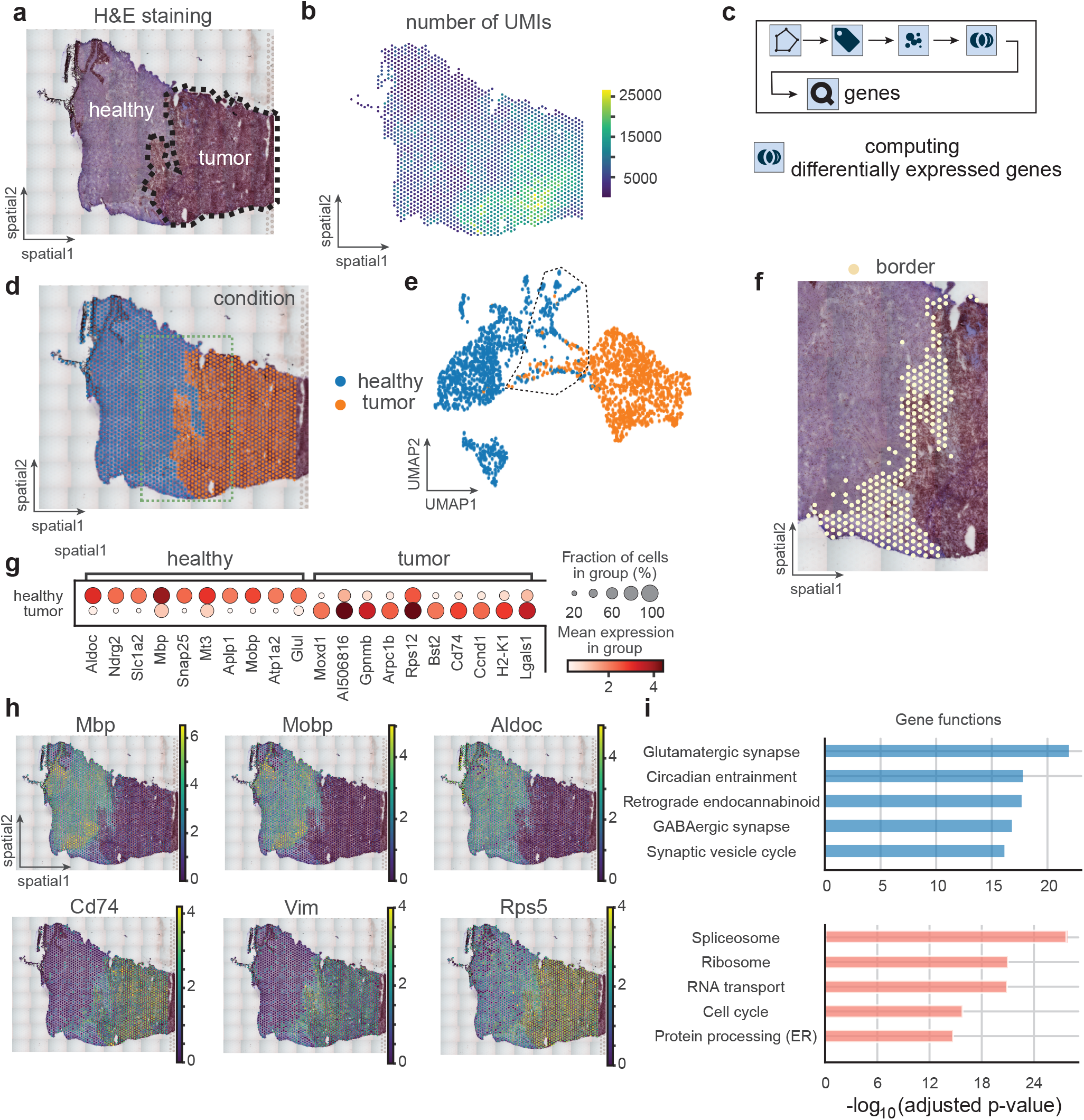
Annotation of a mouse glioblastoma Visium dataset using tissue histology image. **a)** H&E stained image of the tissue region. **b)** the number of transcripts per spot shown in spatial coordinates. **c)** A workflow to annotate spatial transcriptomics datasets. **d)** manual annotation of spots based on H&E image in (a). Blue color represents the healthy region and orange color represents the tumor region. The green dotted line highlights the interface (border) between tumor and healthy regions. **e)** UMAP based on gene expression profiles (transcriptomics) colored by the annotation in (d). The spots between tumor and healthy clusters are outlined by a dotted line. **f)** A zoomed-in version of the H&E image overlaid with spots between the tumor and healthy clusters highlighted by a dotted line in UMAP from (e). **g)** Dot plot showing the top 10 differentially expressed genes in healthy and tumor regions, respectively, by t-test. **h)** Spatial plots for marker gene expression identified from (f). (Top) healthy, and (Bottom) tumor region-specific genes. **i)** Top five pathways from Gene Set Enrichment Analysis (GSEA) from differentially expressed genes identified by t-test in (f). (Top) healthy, and (Bottom) tumor-specific pathways.

In this section, we introduce a vignette using a mouse glioblastoma (brain tumor) dataset acquired with the Visium platform from 10x Genomics. Visium offers sequencing-based transcriptomics covering the whole transcriptome in an unbiased manner. It is, however, limited by the spatial resolution of a unit bearing the same spatial barcodes, called “spot”, whose diameter is 55 μm that typically captures multiple cells with potentially different cell types (Moses and Pachter 2022a). In this sample dataset, there are two distinct regions that correspond to healthy tissue (left, light purple) and the tumor (right, dark purple) (Figure 5a)(Echle et al. 2021; Ru et al. 2023).

Notably, the distinction between healthy and tumor regions is also clear from the sequencing metrics including the number of transcripts per spot, as previously reported (Figure 5b) (Echle et al. 2021; Ru et al. 2023). Based on the overlaid histology image, we annotated the spots as “healthy” or “tumor” using the Cellxgene VIP’s lasso tool (Figure 5c, d). In the gene expression UMAP, the spots are clearly distinguished between healthy and tumor conditions (Figure 5e).

However, some spots appeared scattered between the two clusters, possibly due to a mixture of healthy and tumor cells within a spot, or the presence of intermediate cell states (Figure 5e, outlined by a dotted line). When mapping these spots back to their spatial coordinates, they were indeed localized to the border between healthy and tumor regions (Figure 5f).

Next, we identified genes that are differentially expressed between healthy or tumor regions and checked their spatial gene expression patterns (Figure 5g and h). Gene Set Enrichment Analysis (GSEA) (Fang, Liu, and Peltz 2023) showed enrichment of neuronal pathways in the healthy region, and enrichment of pathways related to cell proliferation (transcription, translation, splicing, transport, etc.) in the tumor region (Figure 5i). Together, we showed a workflow of annotating the spatially resolved transcriptomics data using both spatial and gene expression information complementarily.

## 6. Discussion

We introduced four vignettes to guide scientists through cell-type annotations for single-cell multi-omics datasets of their interest. Interactive software tools such as CZ CELLxGENE facilitate the cell-type annotation procedure with their ability to dynamically switch between embeddings from multiple modalities and utilize multiple modalities jointly. We provided examples of each scenario presented in Figure 1f, where two modalities can provide redundant information (Section 5; Figure 1f, top), one modality can have a superior resolution than the other (Section 3; Figure 1f, center), or two modalities can reveal a hidden population only when both modalities were considered together (Section 4; Figure 1f, bottom). We also showed examples where the degree of complementarity between modalities depends on the cell type of interest.

What is the source of this complementarity between modalities? Overall, the complementarity might stem from two primary sources - technical and biological. First, technical differences exist between the experimental assays of each modality. For example, scRNA-seq is known to have a significant level of dropouts (Kharchenko, Silberstein, and Scadden 2014), whereas CITE-seq’s ADT modality (for surface protein) shows high levels of background protein signal due to non-specific binding of antibodies (Stoeckius et al. 2017; Mulè, Martins, and Tsang 2022).

Second, even with perfect measurements, the information flow along the central dogma might not be consistent for different cellular states, resulting in one modality having more importance when defining these cell types (G.-W. Li and Xie 2011). For example, the immune progenitor subtypes are known to have similar transcriptomic profiles while their epigenetic profiles have distinct characteristics, as chromatin accessibility precedes gene expression (Ma et al. 2020).

In principle, multi-omic technologies can facilitate the identification of heterogeneous populations whose subtle differences were hidden under the lens of uni-modal datasets. We expect that more multi-omics technologies will emerge at the intersection of proteomics and metabolomics, opening new avenues for understanding cellular states at the functional level. We believe that our framework of utilizing complementary information from multiple modalities will be broadly applicable beyond the modalities mentioned in this article.

Lastly, we hope our article will serve as a guideline for biologists, especially those without specific bioinformatics training who seek to analyze their own datasets. The set of tutorials illustrated in this article, together with the accompanying slide decks demonstrating these workflows using the CZ CELLxGENE platform, will equip bench scientists with the tools to interface with their data. This will allow them to explore and annotate cell types by harnessing their domain knowledge in specific biological systems.

## 7. Methods

All Jupyter notebooks used to process the public datasets and generate figures are available in the public Github repository (https://github.com/czbiohub-sf/celltype_annotation_tutorial). We used exCELLxGENE (https://github.com/czbiohub-sf/excellxgene) version 2.9.2 and Cellxgene VIP (https://github.com/interactivereport/cellxgene_VIP) following the installation guidelines.

1. Data preprocessing notes

a. Both single-cell Multiome and CITE-seq datasets are downloaded from GEO (GSE194122). For both single-cell Multiome and CITE-seq datasets, we subsetted for one dataset specified as “site 1 donor 1” using the “batch” key.
b. snRNA-seq: We used only the RNA modality of the single-cell Multiome dataset (“site 1 donor 1”), which is used again in Section 4. The count matrix was log-normalized for gene expression quantification.
c. CITE-seq: The count matrix was already combined between RNA and ADT modalities, and we transformed the RNA counts using log-normalization and ADT counts using CLR-transformation (Centered-Log Ratio), respectively.
d. Single-cell Multiome: The count matrix was already combined between RNA and ATAC modalities.
e. Spatial Transcriptomics: We used a mouse glioblastoma dataset acquired via 10xGenomics’ Visium platform, which captured both tumor and healthy regions of the mouse brain. The dataset was provided by the Genomics platform from Chan Zuckerberg Biohub, SF and Arantxa Tabernero’s lab at the University of Salamanca.

## Resources

1. Tutorial slides are deposited in CZ CELLxGENE’s Documentation page with the following link :

https://cellxgene.cziscience.com/docs/05Annotate%20and%20Analyze%20Your%20Data/5_8Multimodal%20Annotations

## Acknowledgment

We thank the whole Data Science and Genomics platforms of the Chan Zuckerberg Biohub, SF for helpful discussion and support. We are grateful for the support, feedback, and proofreading from Sandra Schmid, Keir Balla, Loic Royer, Leah Dorman, and Merlin Lange. We thank Open Problems in Single Cell consortium for sharing their BMMC single-cell Multiome and CITE-seq datasets which we used for our vignettes. We appreciate the lab of Aranxta Tabernero at the University of Salamanca, especially Aranxta Tabernero, Laura Garcia Vicente, Andrea Álvarez-Vázquez, and Raquel Flores-Hernández for generously sharing their mouse glioblastoma samples and datasets for our spatial transcriptomics section. We also thank the single-cell team at the Chan Zuckerberg Initiative, Jonah Cool, Colin Megil, Ambrose Carr, Max Lombardo, and Kirsty Ewing, for their support and helpful feedback on this work. Funding for this work was provided by Chan Zuckerberg Biohub San Francisco (CZB SF). We thank the CZB SF donors, Priscilla Chan and Mark Zuckerberg, for their generous support

---

**Box 1.**
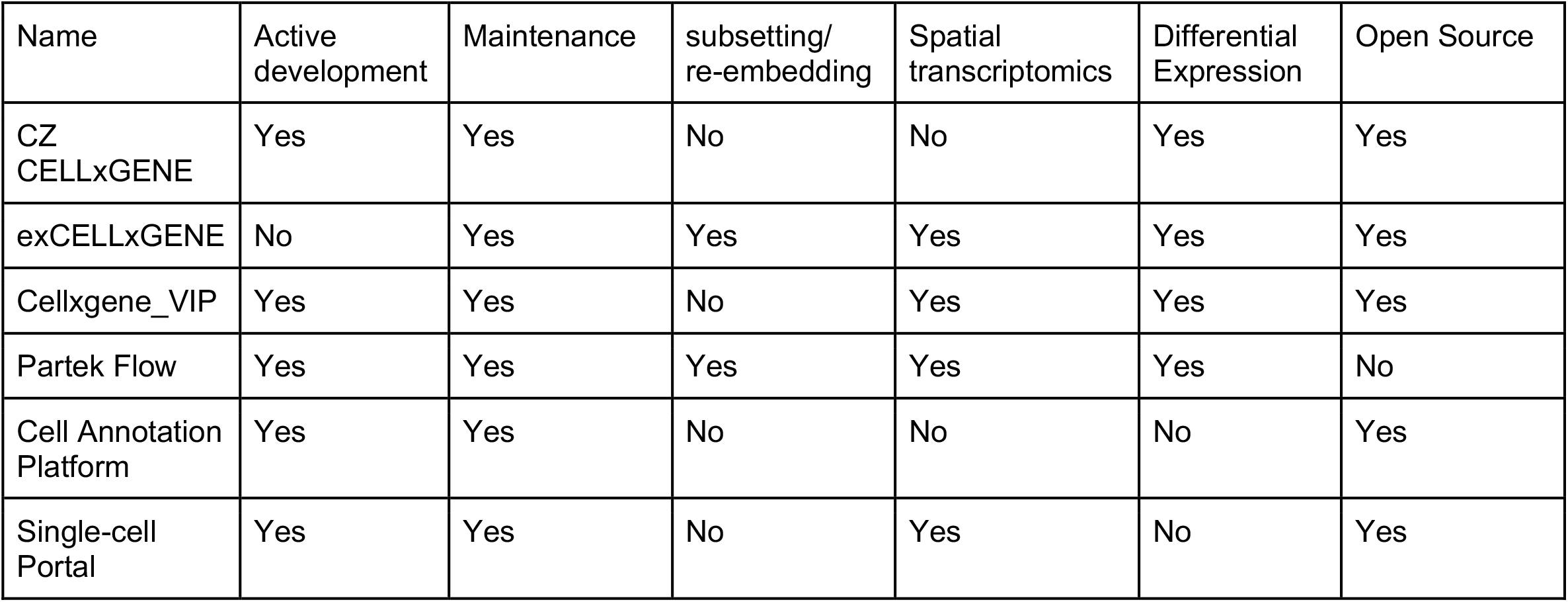
Comparison between interactive software for single-cell genomics data analysis

## Supplementary Material

**Figure S1.**
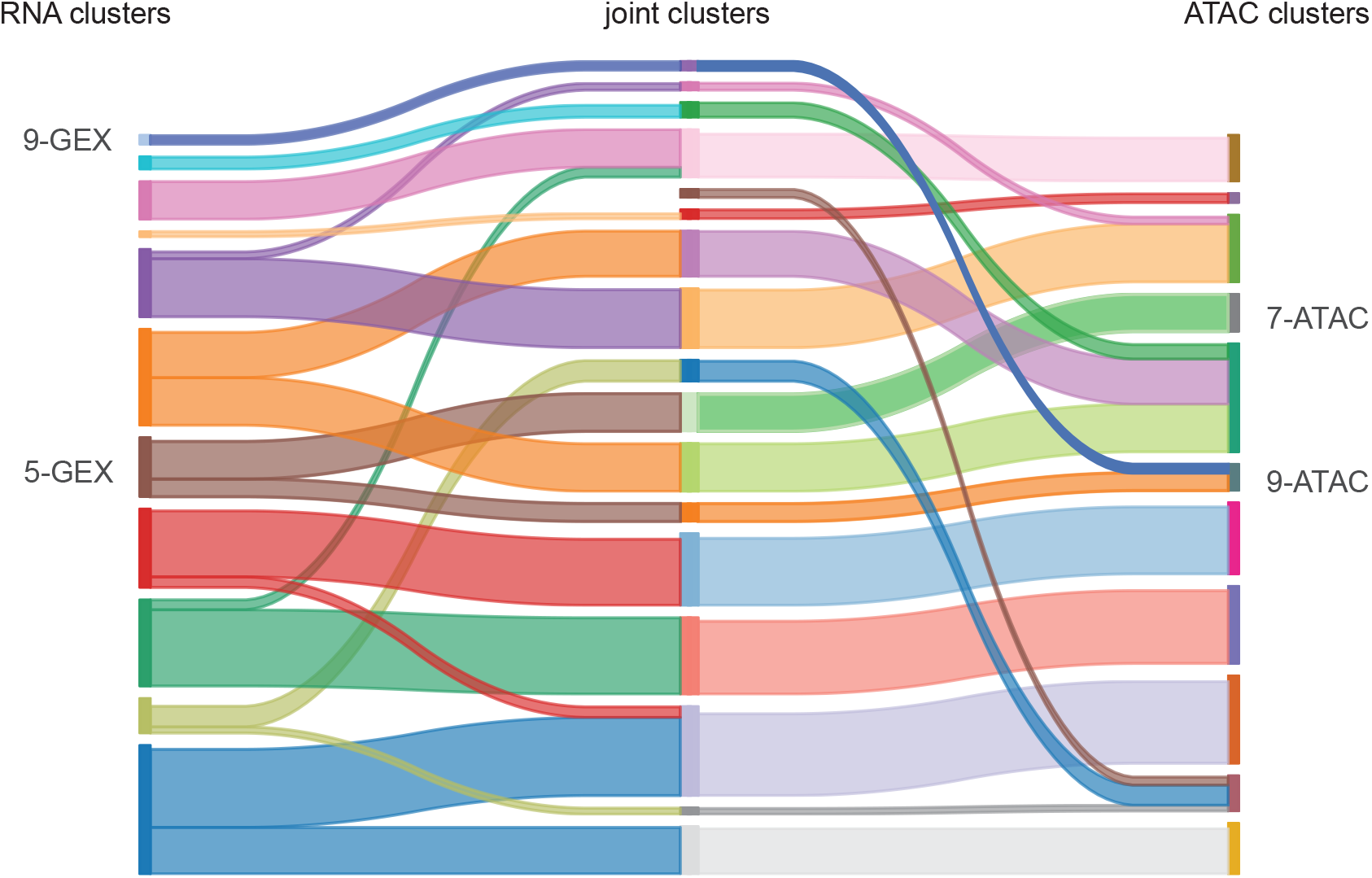
Sankey diagram between Leiden clusters of RNA (Left), ATAC (Right), and joint embeddings (Center) from single-cell Multiome dataset (Figure 4).

## Notes

### Competing Interest Statement

The authors have declared no competing interest.

